# Design of TCR-mimicking binders for pHLA with high potency

**DOI:** 10.1101/2025.10.03.680404

**Authors:** Teng Xie, Yaqi Fu, Shi-Meng Gong, Damiano Burrato, Ruhong Zhou

## Abstract

The rational design of high-specificity binders to peptide–HLA (pHLA) complexes remains a major challenge in personalized immunotherapy, particularly for shared neoantigens with single-point mutations. To address this, we have developed an integrated framework that combines knowledge-based deep learning with physics-based simulation for the design of highly specific pHLA binders. Applied to p53 R175H–HLA-A*02:01, molecular-dynamics–guided electrostatic filtering yielded nanomolar binders with >10-fold selectivity over wild type. Motivated by the challenge of KRAS G12V–HLA-A*03:01, which lacks electrostatic cues, we implemented message-passing neural network (MPNN)–based optimization strategies to broaden binder design beyond charge-dependent interfaces. When reapplied to p53 R175H, these strategies improved success rates from 2/5 to 5/5, with most candidates exhibiting minimal or no binding to the wild-type complex. Chimera-based and residue-retention approaches within the MPNN optimization further expanded the sequence–structure search space while preserving critical hotspot interactions. This integrated framework thus enables the design of nanomolar, mutation-selective binders to pHLA complexes, advancing next-generation personalized cancer immunotherapies.

## Introduction

The development of personalized cancer immunotherapies has been propelled by the identification of tumor-specific neoantigens—8–11 amino acid peptides derived from somatic mutations—presented by HLA-I molecules on tumor cells^1,2^. These neoantigens can elicit potent T-cell responses and are classified as private or shared^3^. Private neoantigens are unique to an individual’s tumor^4^, while shared neoantigens recur across patients due to common oncogenic mutations in proteins such as KRAS, EGFR, TP53, and BRAF^5,6,7^ (Fig. 1a–b). Shared neoantigens, often arising from single-point mutations, provide attractive targets for off-the-shelf therapeutics but pose substantial challenges because of their subtle structural and electrostatic perturbations.

**Fig. 1.**
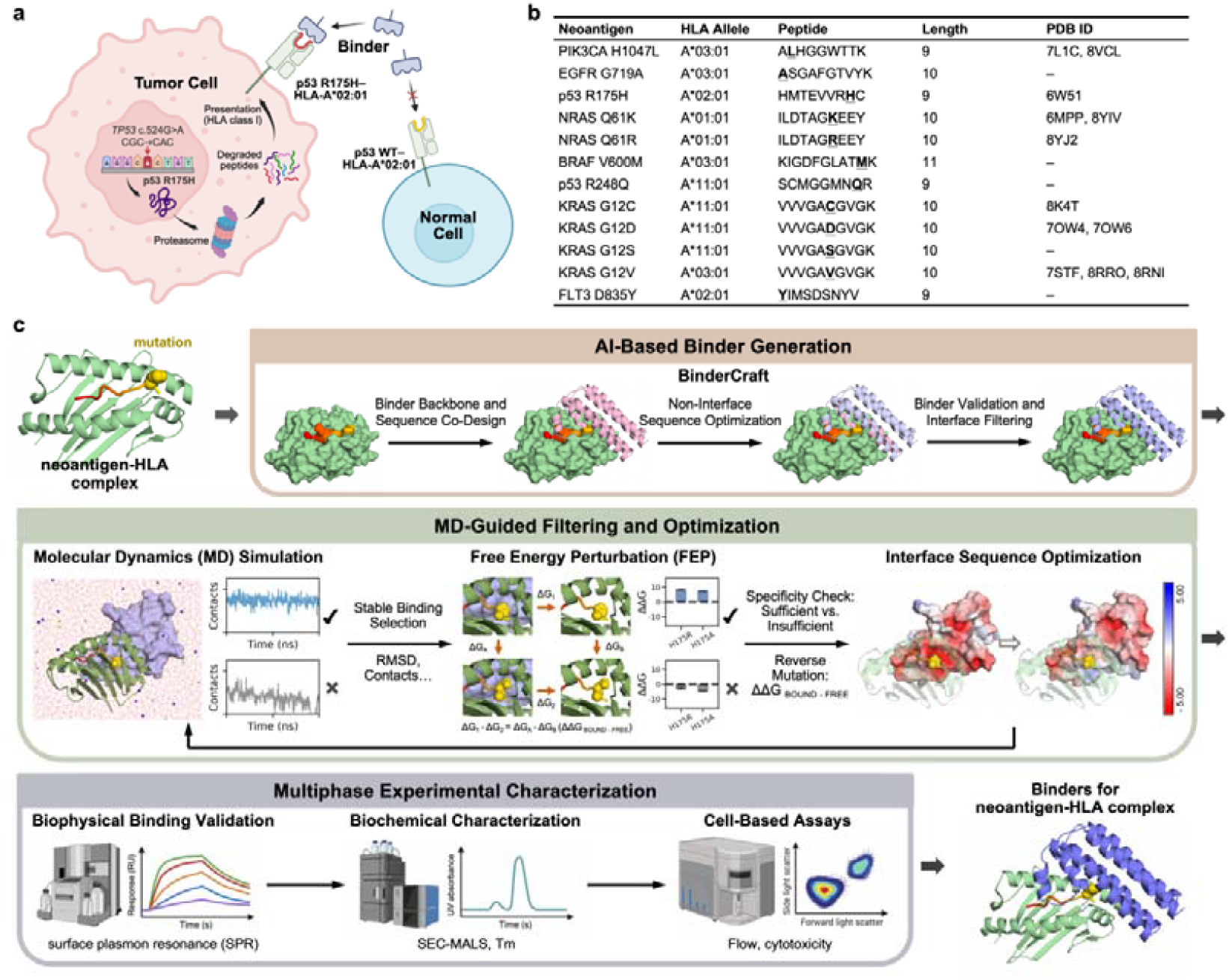
Framework for designing binders targeting single–point-mutated pHLA complexes. **(a)** Schematic illustrating the role of single-mutated neoantigens presented by HLA in cancer cell recognition. **(b)** Table summarizing the reported representative single-mutated pHLA combinations. **(c)** Overview of the framework, integrating AI-based de novo binder generation with physics-based MD-guided filtering and interface optimization, followed by multiphase experimental characterization.

Traditional methods for peptide–HLA (pHLA) binder design rely on protein maturation. Phage display enables high-throughput screening of pHLA antibodies through iterative rounds of binding and amplification, effectively enriching high-affinity binders^8,9^. Yeast surface display combined with computational design facilitates binder optimization by introducing interface diversity prior to experimental selection^10^. Molecular dynamics (MD)-guided refinement was applied to improve antibody specificity by resolving atomic-level interactions at the interface^11^. While powerful, these approaches remain constrained by the requirement for suitable starting scaffolds and limited transferability across mutation contexts.

AI-based strategies, including trained models of RFdiffusion^12^, ProteinMPNN^13^, and AlphaFold2 (AF2)^14^, have recently expanded the toolkit for de novo pHLA binder design. Liu et al. generated small proteins that docked above pMHC peptide-binding grooves and, when displayed on yeast, selectively recognized ten target complexes^15^. Johansen et al. developed a rapid minibinder platform using generative models to design binders targeting the NY-ESO-1 peptide presented by HLA-A*02:01^16^, while Householder et al. created an α-helical TCR mimic achieving nanomolar affinity and strict on-target specificity^17^. These advances highlight the potential of AI-driven approaches to surpass conventional maturation techniques. Other pipelines, such as BindCraft^18^, leverage AF2 through backpropagation to optimize binder sequences; however, AF2 predictions themselves are often insensitive to single-residue changes^19^. Similarly, approximate physics-based scoring functions, widely used in docking and design frameworks, frequently fail to capture the subtle conformational or electrostatic effects of single-point mutations, motivating the use of ensemble methods and rigorous MD-based free energy calculations^20,21,22^.

Building on these principles, we developed an integrated framework that combines knowledge-based deep learning with physics-based simulation for the designing and optimizing binders to single-mutated pHLA complexes (Fig. 1c). Instead of serving only as a validation step, MD–guided refinement was leveraged to resolve local structural rearrangements, electrostatics, and interface complementarity beyond the reach of static scoring. We first applied this framework to p53 R175H–HLA-A*02:01 using electrostatic filtering and refinement, and then extended it to KRAS G12V–HLA-A*03:01 using generalizable sequence–optimization strategies. Reapplying these approaches to p53 R175H confirmed their broader utility, yielding improved success rates compared with electrostatic filtering alone. Together, these advances establish a versatile framework for binder optimization, enabling systematic targeting of cancer-related pHLA.

## Results

### p53 R175H binder interfaces require optimization to achieve specificity

To design binders that selectively recognize p53 R175H (HMTEVVRHC) without binding to the corresponding wild-type (WT) peptide (HMTEVVRRC) presented by HLA-A*02:01, we first applied the de novo binder design pipeline BindCraft to produce candidate backbones and sequences (Fig. 1c). The mutation site, R175H, was designated as the hotspot for interface design. In total, 135 candidate binders were produced, with binding regions positioned around the mutation site. To evaluate their specificity, we implemented an MD–FEP filtering workflow, combining all-atom MD simulations with free energy perturbation (FEP). For each binder–pHLA complex, we quantified the energetic cost of reverting histidine to arginine (H175R). A large positive ΔΔG upon reversal indicated strong discrimination between mutant and WT pHLA. Among the 135 binders, 45 maintained stable binding to p53 R175H–HLA-A*02:01 during 20 ns MD simulations. These 45 complexes were subjected to reverse-mutation FEP, but only a small subset exhibited substantial energetic penalties (ΔΔG), suggesting that most binders lacked strong mutant selectivity.

Interface analysis revealed the basis for this limited discrimination. Strikingly, many binders contained negatively charged residues near R175H. This electrostatic environment also favored the arginine residue present in the WT peptide, which interacted even more strongly with negatively charged surfaces. Indeed, the density of negative charges at the interface correlated with lower ΔΔG values (Fig. 2a–b). Furthermore, in 11 of 45 binder simulations, Na? ions were observed to occupy negatively charged patches near R175H. In addition, ion-binding prediction suggested that other cations, such as Mn^2+^, could also be recruited at these sites (Fig. 2c), consistent with the Mn^2+^-binding site reported in crystal structures^23^. Together, these findings underscore the excessive local charge density at the designed interfaces.

**Fig. 2.**
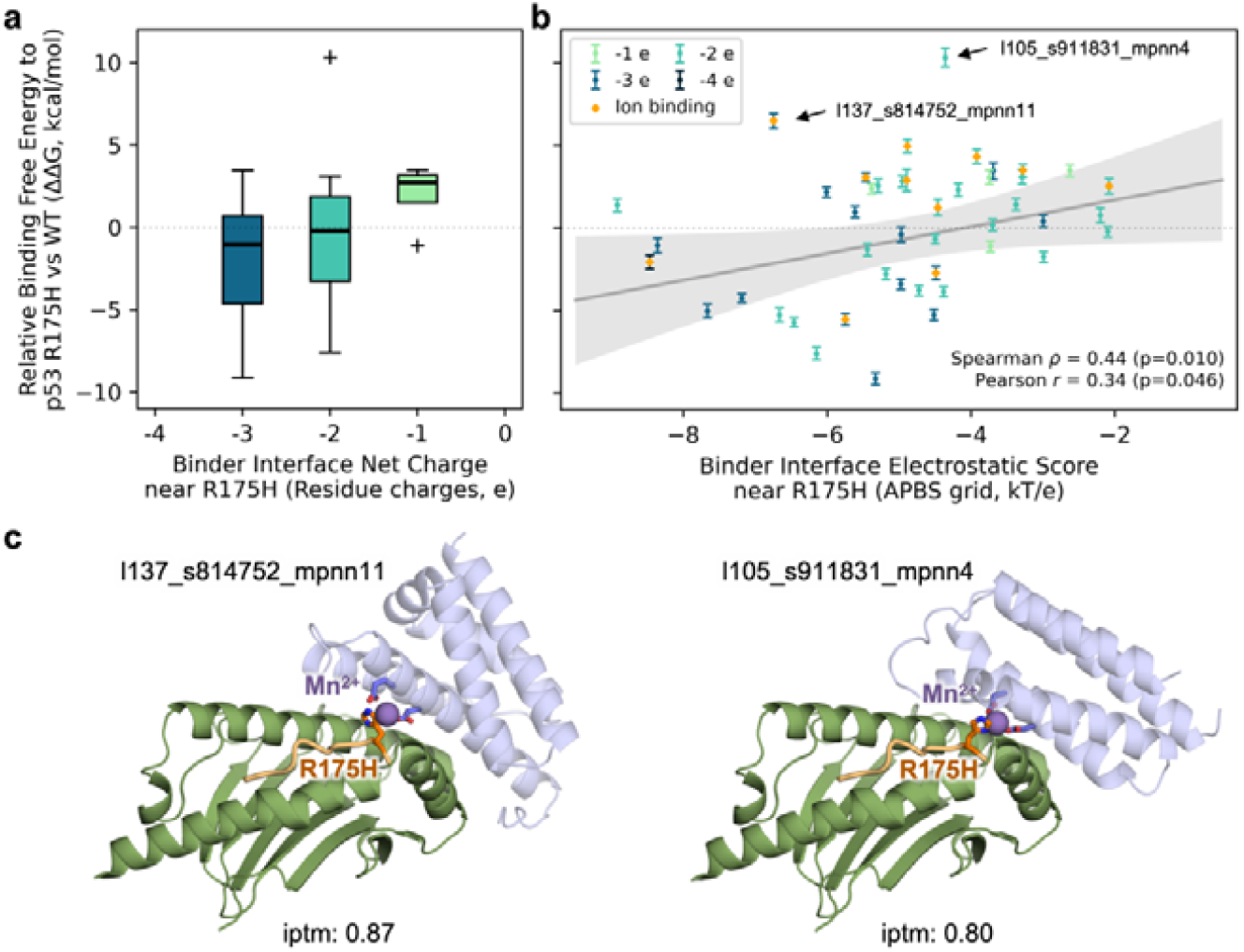
Negatively charged binder interfaces can capture ions and reduce neoantigen specificity. **(a)** Box plot of binding preference (ΔΔG) for binders with varying net charges at the interface near p53 R175H. Binders in which a Na^+^ion was simultaneously coordinated by both R175H and binder residues after MD simulations were excluded from analysis. **(b)** Correlation between the Binder Interface Electrostatic Score near p53 R175H and binding preference (ΔΔG). The score was calculated by averaging APBS-derived electrostatic potential values (see Methods). Scatter points represent individual binder candidates, colored by the net charge of contacting residues (–1, light teal; –4, dark teal). Ion-coordinating binders (orange diamonds) were excluded from the correlation analysis. Correlation outliers (gray hollow circles) represent the 10% of data points with the largest residuals from the linear regression. The gray line indicates the linear regression fit, and the shaded region indicates the 95% confidence interval. **(c)** AlphaFold 3 predictions of two representative binders in complex with p53 R175H–HLA-A*02:01 and Mn^2+^. The Mn^2+^ ion (spheres) coordinates both R175H and negatively charged binder residues (sticks).

Given that our design pipeline relied on AF2 weights (via BindCraft backpropagation through AF2) for interface generation, we hypothesize that the model is biased toward placing negatively charged residues adjacent to histidines. While this may stabilize binding, it fails to discriminate between R175H and WT. These results highlight the need for further optimization of the AI-generated binder interface—particularly through modulation of interfacial charge distribution—to achieve the desired specificity.

### Electrostatic filtering and MD-guided refinement improve p53 R175H binder specificity

To overcome the limitations of the initial AI-generated binders, we developed a multi-step MD-guided optimization workflow. In this pipeline, binders from BinderCraft were first screened for strong contacts with the R175H residue (Step 1). We then applied solMPNN (originally denoted MPNN_sol_)^24^, a ProteinMPNN variant trained on soluble protein assemblies, to mutate residues proximal to R175H while retaining overall interface geometry. Designs that satisfied predefined interface filters^12,18,25,26,27^, along with our electrostatic filters, were advanced (Step 2). Surviving candidates were subjected to MD simulations to assess binding stability and conformational robustness (Step 3). Finally, we applied FEP filtering (Step 4), retaining only those binders with sufficient energetic penalties for reverse (H175R ≥ 6 kcal/mol) and alanine (H175A ≥ 1 kcal/mol) mutations. In a parallel control workflow, interface optimization was replaced by a simplified charge-based selection to assess the specific contribution of solMPNN-based refinement.

This workflow yielded a high success rate: the majority of solMPNN-refined binders (11/15) passed the MD filters, whereas none of the control binders (0/1) did. FEP calculations further confirmed that a subset of refined binders (5/11) strongly discriminated against p53 WT, meeting both ΔΔG thresholds (Fig. 3a). Sequence analysis revealed that solMPNN refinement reduced excessive negative charge and favored a balanced distribution of hydrophobic and polar residues at the interface (Fig. 3b). Notably, binders that passed FEP filtering converged on recurrent hydrophobic and polar substitutions adjacent to R175H, underscoring the importance of charge rebalancing for specificity.

**Fig. 3.**
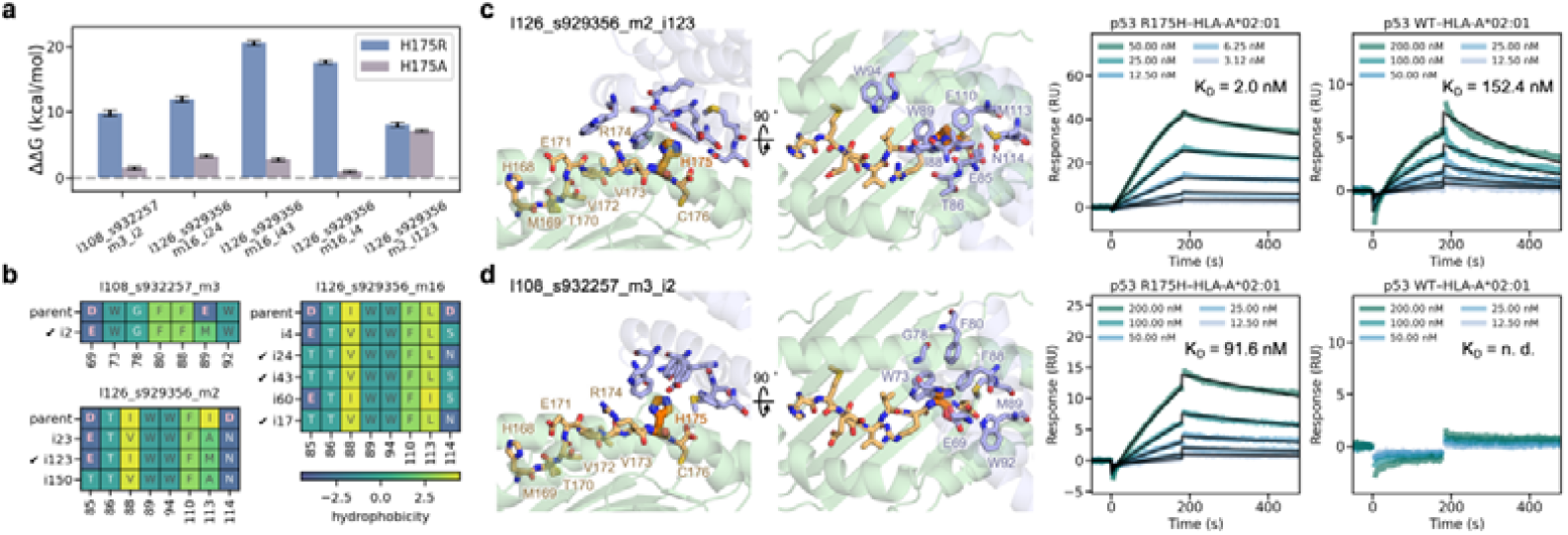
Electrostatic filtering and MD-guided interface refinement improve specificity of p53 R175H–HLA-A*02:01 binders. **(a)** FEP results for reverse (H175R) and alanine (H175A) mutations of interface-optimized binders that passed FEP filtering. Binder IDs use the following notation: l = length, s = seed, m = MPNN model, i = interface-optimized sequence. **(b)** Interface sequence alignment of binders passing FEP filters compared with their parent and sibling binders (sequence variants derived from the same parent binder through different mutation paths). Residue boxes are colored by GRAVY hydrophobicity (see colorbar), with residue letters colored by type (“negative”: pink; “polar”: light gray; “hydrophobic”: dark gray). SPR-validated candidates are marked with ticks. **(c-d)** Structural models and SPR data for two confirmed binders. Neoantigens (orange) and binder interface residues (lignt blue) are shown as sticks. SPR sensorgrams show binding to both p53 R175H and p53 WT–HLA-A*02:01.

To experimentally validate these predictions, we tested all FEP-passing binders using surface plasmon resonance (SPR) (Supplementary Table 1). Two candidates displayed high-affinity binding to p53 R175H–HLA-A*02:01, with dissociation constants (*K*_d_) of 2.0 nM and 91.6 nM, while exhibiting weaker binding (152.4 nM) or no detectable interaction with the WT complex (Fig. 3d–e). Structural models revealed that their optimized interfaces formed stabilizing interactions with the mutant histidine side chain, consistent with selective recognition of the R175H mutation. These findings demonstrate that MD-guided interface refinement enhances the specificity of AI-generated binders and enables the discovery of candidates with experimentally validated neoantigen selectivity.

### General MPNN-based optimization strategies enhance KRAS G12V binder interfaces

To test whether our workflow could be generalized to other pHLA systems, we selected KRAS G12V (VVVGAVGVGK) presented by HLA-A*03:01, a single-point mutation with distinct structural and physicochemical features, and applied BinderCraft to generate initial binders. Following preliminary screening for contacts with the G12V hotspot and subsequent MD–FEP filtering, only a small subset of candidates exhibited sufficient stability and specificity, defined as a reverse-mutation penalty (V12G ΔΔG) ≥ 6 kcal/mol. To improve their performance, we incorporated five complementary MPNN-based strategies with predefined interface filters: solMPNN (S1), solMPNN with residue retention (S2), solMPNN chimera (S3), solMPNN chimera with residue retention (S4), and ThermoMPNN^28^ chimera (S5). In chimera-based approaches, the pHLA and binder were physically linked with a flexible linker and treated as a single polypeptide chain to enable direct optimization of interfacial residues. In residue-retention approaches, stable residues proximal to G12V were preserved to maintain favorable contacts. These optimization schemes were selectively applied to 9 binders showing limited—but non-negligible—discrimination against the reverse mutation (V12G ΔΔG ≥ 3 kcal/mol), and the resulting variants were re-evaluated by MD and FEP filtering. As a control, solMPNN was applied to binders that failed initial MD filtering, providing a direct benchmark for assessing the benefit of optimizing promising candidates.

After the five strategies, more than half of the promising candidates (5/9) yielded mutants that successfully passed both MD and FEP filtering, though with substantial differences in efficiency (Fig. 4a and Supplementary Table 2). solMPNN chimera with residue retention (S4) and ThermoMPNN chimera (S5) consistently generated the highest number of viable binders, suggesting that chimera-based approaches and residue-retention strategies substantially improve interface design. Analysis of interface hydrophobicity and MD stability revealed that MPNN-optimized binders shifted toward a more favorable distribution, characterized by increased binder hydrophobicity at the mutant site coupled with improved contacting atoms (Fig. 4b–c and Supplementary Fig. 1). By contrast, the control workflow rarely rescued unstable binders, underscoring the importance of applying MPNN-based optimization to promising starting points.

**Fig. 4.**
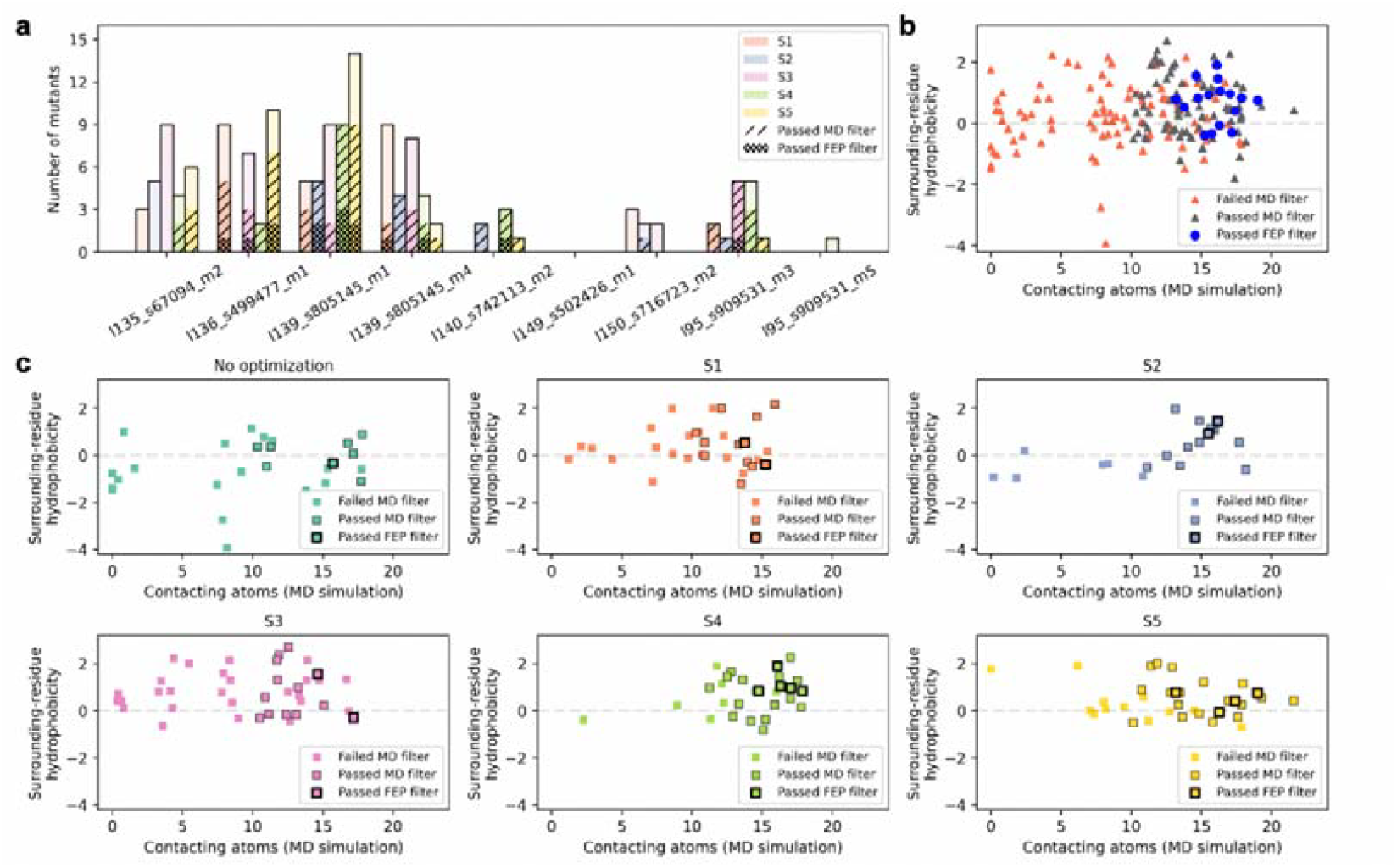
General MPNN-based optimization strategies enhance KRAS G12V–HLA-A*03:01 binder interfaces. **(a)** Number of mutants after each MPNN-based strategy (S1–S5), with those passing MD and FEP filtering highlighted. **(b)** Interface hydrophobicity vs. MD stability (contacting atoms with G12V) for all binders, with MD- and FEP-passing binders highlighted. **(c)** Interface hydrophobicity vs. MD stability before and after optimization by each strategy (S1–S5), with MD- and FEP-passing binders highlighted.

Together, these results demonstrate that integrating diverse MPNN-based strategies enhances binder interface optimization for KRAS G12V. In particular, approaches that balance residue retention or thermodynamic constraints proved most effective, supporting a generalizable framework that unites physics-based filtering with generative optimization.

### General MPNN-based optimization strategies extend to p53 R175H binders and outperform electrostatic filtering

We next applied our generalizable workflow to optimize binders targeting p53 R175H presented by HLA-A*02:01, allowing direct comparison with the charge-balanced optimization strategy previously applied to the same system (Fig. 3). From the same 135 initial BinderCraft designs, binders that established moderate contacts with R175H were subjected to MD and FEP filtering, which yielded no passing candidates but identified nine promising designs with measurable, though modest, discrimination against the reverse mutation (H175R ΔΔG ≥ 3 kcal/mol). To improve these binders, we applied three MPNN-based strategies — solMPNN chimera (S3), solMPNN chimera with residue retention (S4), and ThermoMPNN chimera (S5) — followed by a second round of MD and FEP filtering.

As observed for KRAS G12V, all nine promising R175H candidates yielded mutants that successfully passed both MD and FEP filtering after MPNN-guided optimization, though the efficiency of each strategy varied across candidates (Fig. 5a). In contrast to the KRAS G12V system, however, R175H-optimized binders did not exhibit a concentrated shift toward increased interface hydrophobicity and enhanced contact profiles (Fig. 5b), likely reflecting distinct physicochemical requirements of valine versus histidine recognition. Sequence-level comparisons of two promising candidates revealed that successful binders converged on more hydrophobic and less negatively charged substitutions at key contact positions, while maintaining favorable polar interactions (Fig. 5c and Supplementary Fig. 2).

**Fig. 5.**
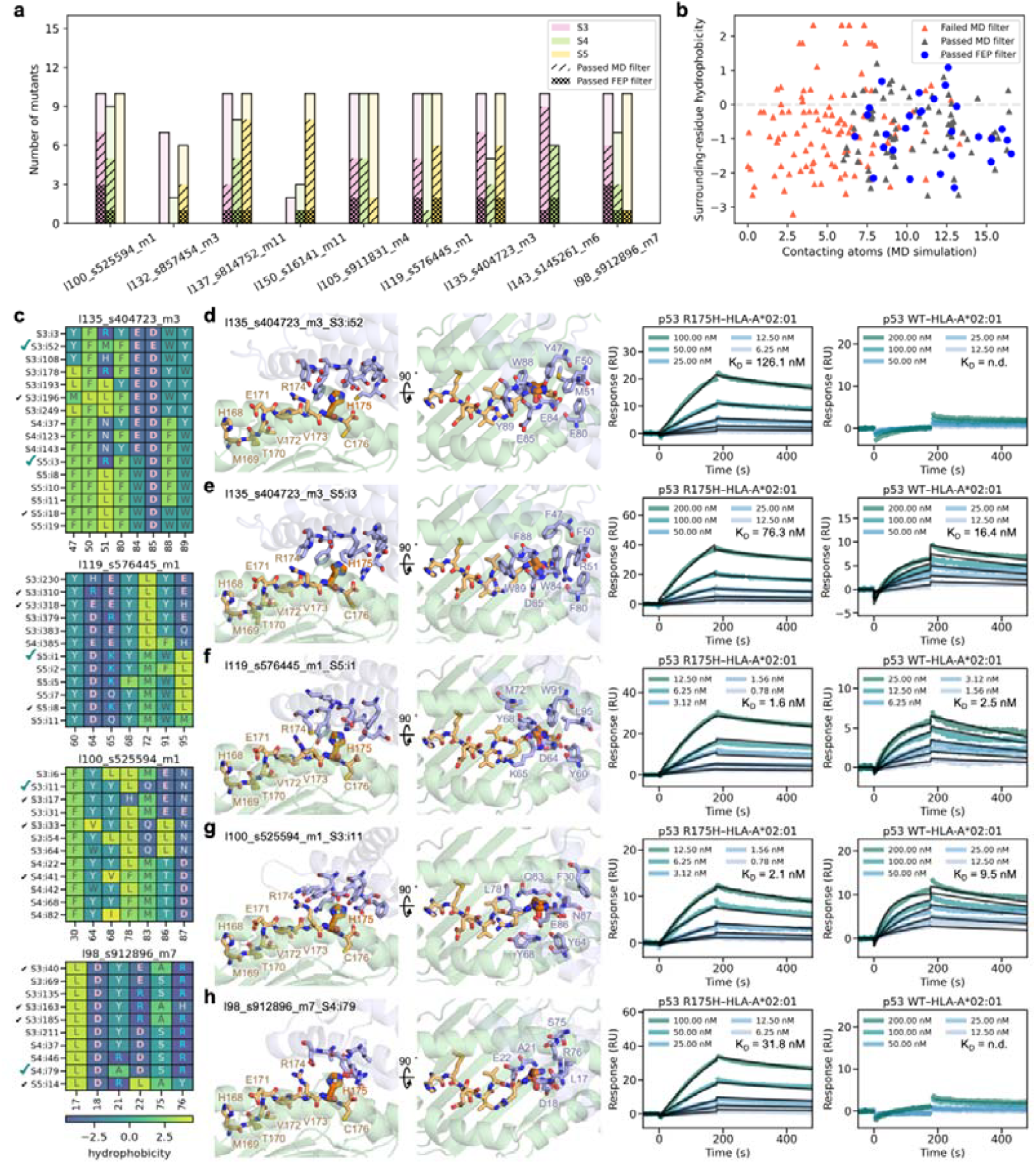
General MPNN-based optimization strategies extend to p53 R175H–HLA-A*02:01 binders. **(a)** Number of mutants after each MPNN-based strategy (S3–S5), with those passing MD and FEP filtering highlighted. **(b)** Interface hydrophobicity versus MD stability (atoms contacting R175H) for all binders, with MD- and FEP-passing binders highlighted. **(c)** Interface sequence alignment of MD-passing binders compared with their parent. Residue boxes are colored by GRAVY hydrophobicity (see colorbar), with residue letters colored by type (negative: pink; positive: cyan; polar: light gray; hydrophobic: dark gray). FEP-passing candidates are marked with black ticks; SPR-validated binders with teal ticks. Binder IDs use the following notation: l = length, s = seed, m = MPNN model, S = strategy, i = interface-optimized sequence. **(d-h)** Structural models and SPR data for five confirmed binders. Neoantigens (orange) and binder interface residues (lignt blue) are shown as sticks. SPR sensorgrams show binding to both p53 R175H and p53 WT–HLA-A*02:01.

Five mutants derived from four promising candidates were selected for SPR validation (Supplementary Table 3), all showing selective recognition of p53 R175H over p53 WT– HLA-A*02:01 (Fig. 5d–f). All five binders bound p53 R175H–HLA-A*02:01 with nanomolar affinity (*K*_d_ = 1.6–126.1 nM). Two binders exhibited weaker binding to the WT complex (1.6 nM for R175H vs. 2.5 nM for WT; 2.1 nM for R175H vs. 9.5 nM for WT), two showed no detectable WT binding (126.1 nM and 31.8 nM for R175H), and one displayed stronger binding to WT than to R175H (76.3 nM vs. 16.4 nM). Interestingly, the stronger-binding WT binder and the non-binding WT binder originated from the same promising candidate, highlighting the context-dependence of interface specificity. Collectively, these results validate the predictive accuracy and efficiency of our integrated workflow and demonstrate its ability to generate experimentally verified, neoantigen-specific binders across distinct mutation contexts.

## Discussion

Our results highlight the critical importance of precise optimization of pHLA binder interfaces when targeting single-point mutations such as p53 R175H and KRAS G12V. Initial AI-generated binders often fail to discriminate between mutant and wild-type alleles due to subtle electrostatic or conformational differences and limitations of AF2’s sensitivity. Interfacial mutation and filtering combined with MD- and FEP-guided selection corrected these biases, improving specificity by mitigating unfavorable charge distributions and interface dynamics. FEP served as a final screen to estimate mutation-induced energy changes for most scenarios. Notably, one p53 R175H binder displayed stronger binding to WT than R175H (76.3 nM vs. 16.4 nM), highlighting FEP’s limitations in capturing subtle conformational or ion-mediated effects and underscoring the need for experimental validation.

To extend optimization, we applied general MPNN-based strategies. SolMPNN and thermoMPNN, while not specifically trained on chimeric binder–pHLA complexes, capture broad sequence–structure and stability determinants that can enhance interfacial complementarity. Chimera strategies further exploit evolutionary-like information embedded in protein fold space. A similar continuity is seen with AlphaFold2 and AlphaFold-Multimer^29,30^, where models trained on monomers can accurately predict complexes. Guided by this rationale, MPNN- and chimera-based optimization improved KRAS G12V binder selectivity and outperformed electrostatic filtering for p53 R175H (binding success rate: 2/5 → 5/5; specificity success rate: 1/5 → 2/5), demonstrating transferable principles of interface design across distinct neoantigen contexts. Retaining stable interface residues during MD simulations often facilitated further optimization, highlighting the value of integrating physics-based ensemble information with AI-driven strategies. Employing multiple MPNN-based approaches broadens accessible fold space and mitigates potential constraints, pointing to the need for future models that better integrate physical and evolutionary principles.

Structural comparison of our binders with antibody H2 for p53 R175H–HLA-A*02:01 (Supplementary Fig. 3) revealed distinct interface engagement modes. Our binders primarily employ α-helical motifs, whereas H2 uses loop-mediated contacts. Consequently, our binders generally make fewer direct contacts with R175H-distal residues, yet binding affinities and mutant-specific recognition are comparable. This indicates that different structural scaffolds can achieve similar functional outcomes, though divergent specificity toward other pHLA complexes may arise, highlighting both opportunities and challenges in designing neoantigen-specific binders with alternative motifs.

Beyond these specific findings, our study points to broader lessons for AI-driven design. AlphaFold2 has been reported to recover metalloprotein motifs such as Fe–S clusters and Zn-binding sites^31^, and AF2-derived models can introduce systematic biases, such as spurious ion-binding motifs, underscoring the need for physics-based correction. De novo scaffolds remain essential starting points that, once filtered by MD and FEP, can yield high-affinity and specific binders, supporting prior findings that appropriate topologies coupled with sequence diversification are key to achieving high-affinity and specificity^32^. Finally, distal mutations can influence specificity through long-range effects (see KRAS G12V interface in Supplementary Fig. 1), highlighting the need to consider not only direct contacts but also allosteric contributions^33,34^.

In summary, we present an integrated framework that combines knowledge-based deep learning with physics-based simulation for optimizing binders against single-mutated pHLA complexes. By combining AI-guided diversification with physics-based refinement, this framework not only enhances the specificity for targeting the very challenging single-point mutations, but also provides a versatile platform to accelerate the development of personalized cancer immunotherapies.

## Methods

### pHLA binder generation

Target models for de novo binder design were constructed from truncated HLA-A (residues 1–180) structures and their associated neoantigen peptides obtained from the Protein Data Bank: p53 R175H–HLA-A*02:01 (PDB ID: 6W51) and KRAS G12V– HLA*A03:01 (PDB ID: 7STF). Each HLA–peptide complex was linked via a flexible GGGGSGGGGSGGGG linker to form a single polypeptide chain suitable for binder design and subsequent optimization. To avoid steric clashes with the linker, the N-terminal residue of each neoantigen was replaced with alanine. Both alanine substitutions and linker modeling were performed using PyRosetta^35^. The resulting structures served as input PDBs for BinderCraft^18^ (“design_algorithm”: “4stage”; “predict_initial_guess”: true). Binder lengths were set to 80–150 residues, and hotspot residues were defined as R175H (residue 202 in p53 R175H–linker–HLA-A*02:01) and G12V (residue 200 in KRAS G12V–linker–HLA-A*02:01). Binder lengths were set to range from 80–150 residues. Generated binders were filtered with the default BinderCraft interface filters to select high-quality candidates for subsequent optimization.

### MPNN-based interface optimization

Interface optimization was performed using MPNN-based strategies. SolMPNN and ThermoMPNN were applied to introduce mutations at non-interface residues as well as interface residues within 6 Å of the hotspot (R175H or G12V). More distal interface residues were retained to preserve baseline binding interactions. To possible long-range influences, mutant variants sharing identical hotspot-proximal residues but differing in non-interface positions were selected for further evaluation.

For the electrostatic filtering strategy applied to p53 R175H binders, solMPNN-generated mutants were subjected to predefined interface filters in BinderCraft, and further selected if they met one of the following criteria: (i) residues contacting R175H (<6 Å) with a net charge between 0 and 1 and ≥16 binder atoms contacting R175H (<4 Å), or (ii) residues contacting R175H (<6 Å) with a net charge ≤0 and ≥6 binder atoms contacting R175H (<4 Å). Redundant binders were removed.

For the five general MPNN-based optimization strategies — solMPNN (S1), solMPNN with residue retention (S2), solMPNN chimera (S3), solMPNN chimera with residue retention (S4), and thermoMPNN chimera (S5) — applied to both p53 R175H and KRAS G12V binders, mutants were retained if they passed BindCraft filters and contained ≥6 (p53 R175H) or ≥16 (KRAS G12V) binder atoms in contact with the hotspot residue within 4 Å. In chimera-based approaches, the pHLA and binder were physically linked with a flexible (GGGGS)n linker and treated as a single polypeptide chain to enable direct optimization of interfacial residues. The initial linker length was determined by measuring the distance between the binder C-terminus and the HLA–linker–neoantigen N-terminus, using the relation:

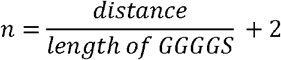

If the linker was insufficiently long, n was incremented until a suitable length was achieved.

In residue-retention strategies, residues forming stable hydrogen bonds or exhibiting low fluctuations within 6 Å of the hotspot were preserved to maintain favorable contacts during optimization.

### Molecular dynamics simulations

Candidate binder–pHLA complexes were selected for all-atom MD simulations. Alanine substitutions and flexible linkers in HLA–linker–neoantigen chimeras were remodelled to recover the original pHLA complex using VMD^36^ scripts. Native disulfide bonds in HLA were added where appropriate. All systems were solvated with TIP3P water and neutralized with 0.15 M KCl. Final system sizes ranged from ∼70,000 to 120,000 atoms, depending on the size of binders.

Simulations were performed with GROMACS version 2020.6^37,38^ using CHARMM36m protein force field. Long-range electrostatics were treated with the particle mesh Ewald method^39^, and Lennard–Jones interactions were computed with a 12 Å cutoff and a switching function starting at 10 Å. Systems were maintained at 310 K using the V-rescale thermostatL^40^ with a coupling constant of 1 ps, and 1 bar using the Parrinello-Rahman barostat^41^ with a coupling constant of 5 ps. All bonds involving hydrogen atoms were constrained using the LINCS algorithm^42^, allowing for a 2Lfs integration time step. Prior to production, each system was energy-minimized and equilibrated in both the canonical (NVT) and isothermal–isobaric (NPT) ensembles. During equilibration, temperature was ramped to 310 K and positional restraints (1000 kJ/mol·nm^2^) on protein heavy atoms were gradually released (side chains followed by non-Cα atoms). Production simulations were then performed for 200Lns.

### Analysis of the binder interface

Collective variables (CVs) were used to characterize the interface in the predicted binder–pHLA complexes and corresponding MD trajectories. A Binder Interface Electrostatic Score was computed using APBS-derived potential grids (units: kT/e). For each binder structure, all grid points surrounding the p53 R175H Cβ atom were identified, and their potential values averaged, providing a grid-spacing-independent measure of the local electrostatic potential at the binder–R175H interface.

Binder stability and interface interactions were analyzed using VMD^36^. RMSDs were calculated for non-hydrogen atoms of the neoantigen and binder. Contacts were defined as atom pairs within 4 Å between hotspot residues (R175H or G12V) and the binder. Hydrogen bonds were identified using a donor–acceptor distance cutoff of 3.5 Å and an angle cutoff of 40°. Local hydration was assessed by quantifying water molecules within 4 Å of hotspot residues.

### Free energy calculation

The energetic contributions of reverse and alanine mutations of neoantigens were quantified using free energy perturbation combined with Hamiltonian replica-exchange molecular dynamics (FEP/HREX) for improved convergence efficiency^43,44,45,46^. Binding free energy differences (ΔΔG) were estimated by calculating the free energy of residue annihilation in two states: the bound state (Δ*G*_*A*_ ; pHLA bound to the binder) and the unbound state (Δ*G*_*B*_ ; free pHLA). The net free energy contribution of a mutation was obtained via a thermodynamic cycle:

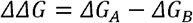

FEP/HREX simulations were performed in GROMACS 2020.6 using the CHARMM36 force field and system topology files generated by PMX^47,48^. Each mutation was sampled across 36 λ-windows, progressively decoupling electrostatic and van der Waals interactions. Simulations were initiated from MD-derived conformations in which key collective variables (RMSD and contacting atoms) were stable.

Every annihilation calculation was run for at least 94 ns (1.3 ns × 36 windowsL×L2 states, with the unbound state shared across pHLAs). Soft-core potentials^49^ were applied (α = 0.5, λ-power = 1, radial-power = 6, and σ = 0.3) to avoid singularities during decoupling. Replica exchanges between adjacent windows were attempted every 1 ps, and Hamiltonians were recorded every 0.2 ps.

Free energy differences (ΔΔG) and associated statistical uncertainties were estimated from the final 1 ns of each window using the Multistate Bennett Acceptance Ratio (MBAR) method^50^, as implemented in Alchemical Analysis^51^.

### Surface plasmon resonance (SPR)

Binder affinities were measured on a Biacore X100 instrument (Cytiva) using multi-cycle kinetics with an SA chip. Biotinylated p53 R175H–HLA-A*02:01 or p53 WT–HLA-A*02:01 complexes were immobilized according to the manufacturer’s instructions. Binders were injected as analytes at defined concentrations in HEPES buffer (pH 7.4) under multi-cycle kinetics mode. Each cycle consisted of a 180 s association phase followed by a 300 s dissociation phase. Data were acquired using Biacore X100 system control software (version 2.0.1.201), and kinetic parameters (*K*_d_, *k*_on_ and *k*_off_) were determined with BIAevaluation software (version 2.0.1.201).

For comparison, the p53 R175H–HLA-A*02:01 antibody H2 was used as a control. Hsiue et al. measured H2 binding using single-cycle kinetics on an SA chip with biotinylated pHLA immobilized. The bispecific antibody was engineered for effective 1:1 binding stoichiometry, enabling fitting with a standard Langmuir 1:1 binding model. Because both datasets employed the same ligand orientation (biotinylated pHLA on SA) and were analyzed with 1:1 kinetic models, the apparent *K*_d_ are directly comparable, despite the different kinetic modes (multi-cycle versus single-cycle). For transparency, *k*_on_ and *k*_off_ values for binder measurements are reported in Supplementary Fig. 3, noting that minor assay-dependent differences may influence comparisons.

## Acknowledgments

This work was partially supported by the National Key R&D Program of China (2024YFA1306400, 2021YFA1201200, 2024YFA1307500), the National Natural Science Foundation of China (U1967217), the National Center of Technology Innovation for Biopharmaceuticals (NCTIB2022HS02010), Shanghai Artificial Intelligence Lab (P22KN00272), the National Independent Innovation Demonstration Zone Shanghai Zhangjiang Major Projects (ZJZX2020014), the Starry Night Science Fund of Zhejiang University Shanghai Institute for Advanced Study (SN-ZJU-SIAS-003), and Zhejiang University Global Partnership Fund (188170+194452505).

---

**Supplementary Table 1.** p53 R175H–HLA-A*02:01 binder candidates from electrostatic filtering that passed FEP filtering.

| Rank_score | Interface optimization | Binder_candidate | Reverse_FEP (ΔΔG, kcal/mol) | Alanine_FEP (ΔΔG, kcal/mol) | Average_i_pTM | Average_i_pAE |
| --- | --- | --- | --- | --- | --- | --- |
| 0.87 | Electrostatic filtering | l126_s929356_m16_i17_r1 | 26.29 | 5.26 | 0.81 | 0.23 |
| 0.79 | Electrostatic filtering | l126_s929356_m16_i43_r3 | 20.51 | 2.78 | 0.80 | 0.24 |
| 0.73 | Electrostatic filtering | l126_s929356_m16_i24_r2 | 11.96 | 3.31 | 0.85 | 0.2 |
| 0.69 | Electrostatic filtering | l126_s929356_m2_i123_r4 | 8.05 | 7.12 | 0.85 | 0.2 |
| 0.61 | Electrostatic filtering | l108_s932257_m3_i2_r4 | 9.82 | 1.50 | 0.71 | 0.32 |

"Rank score" = 0.7 × "Average\_i\_pTM" + 0.3 × "Reverse\_FEP" / "Reverse\_FEP\_max";

**Supplementary Table 2.** KRAS G12V–HLA-A*03:01 binder candidates from general MPNN-based optimization strategies that passed FEP filtering.

| Rank_score | MPNN-based strategy | Binder_candidate | Reverse_FEP ($\Delta\Delta G$, kcal/mol) | Alanine_FEP ($\Delta\Delta G$, kcal/mol) | Average_i_pTM | Average_i_pAE |
| --- | --- | --- | --- | --- | --- | --- |
| **0.89** | **S4** | **I139_s805145_m4_i9_r1** | **7.8** | **3.47** | **0.84** | **0.21** |
| 0.86 | S1 | I139_s805145_m4_i7_r1 | 7.07 | 3.43 | 0.84 | 0.22 |
| **0.85** | **S4** | **I139_s805145_m1_i8_r2** | **6.73** | **3.36** | **0.84** | **0.21** |
| 0.84 | S4 | I139_s805145_m1_i19_r2 | 6.49 | 2.53 | 0.85 | 0.2 |
| 0.84 | S5 | I139_s805145_m1_i2_r1 | 6.67 | 1.86 | 0.84 | 0.22 |
| 0.84 | S5 | I136_s499477_m1_i60_r1 | 7.26 | 2.66 | 0.8 | 0.23 |
| 0.83 | S4 | I139_s805145_m1_i20_r2 | 5.95 | 3.24 | 0.86 | 0.2 |
| 0.82 | S5 | I139_s805145_m1_i10_r2 | 5.96 | 1.23 | 0.85 | 0.21 |
| **0.82** | **S5** | **I136_s499477_m1_i80_r2** | **6.68** | **3.19** | **0.81** | **0.23** |
| 0.82 | S2 | I139_s805145_m1_i7_r1 | 5.97 | 3 | 0.84 | 0.21 |
| 0.81 | Parent | I135_s67094_m2 | 6.38 | 3.06 | 0.8 | 0.25 |
| **0.80** | **S3** | **I136_s499477_m1_i39_r1** | **6.05** | **2.73** | **0.81** | **0.23** |
| 0.80 | S2 | I139_s805145_m1_i9_r1 | 6.14 | 2.75 | 0.8 | 0.26 |
| 0.78 | S1 | I136_s499477_m1_i8_r1 | 5.98 | 2.76 | 0.78 | 0.27 |
| 0.77 | S4 | I140_s742113_m2_i94_r1 | 6.67 | 2.85 | 0.74 | 0.29 |
| 0.74 | S3 | I95_s909531_m3_i71_r1 | 6.08 | 3.29 | 0.72 | 0.3 |

Binder IDs use the following notation: l = length, s = seed, m = MPNN sequence, S = strategy, i = interface-optimized sequence, r = run of interface optimization. Binders selected for SPR validation are highlighted in bold.

**Supplementary Table 3.** p53 R175H–HLA-A*02:01 binder candidates from general MPNN-based optimization strategies that passed FEP filtering.

| Rank_score | MPNN-based strategy | Binder_candidate | Reverse_FEP ($\Delta\Delta G$, kcal/mol) | Alanine_FEP ($\Delta\Delta G$, kcal/mol) | Average_i_pTM | Average_i_pAE |
| --- | --- | --- | --- | --- | --- | --- |
| 0.77 | S3 | l135_s404723_m3_i52 | 12.65 | 1.28 | 0.90 | 0.17 |
| 0.77 | S5 | l135_s404723_m3_i3 | 13.95 | 2.80 | 0.87 | 0.18 |
| 0.73 | S5 | l119_s576445_m1_i1 | 14.8 | 2.38 | 0.80 | 0.24 |
| 0.72 | S4 | l98_s912896_m7_i79 | 8.90 | 1.49 | 0.89 | 0.17 |
| 0.72 | S3 | l100_s525594_m1_i11 | 9.15 | 2.58 | 0.88 | 0.18 |

Binder IDs use the following notation: l = length, s = seed, m = MPNN sequence, S = strategy, i = interface-optimized sequence. The interface optimization run number was set to 1.

**Supplementary Fig 1.**
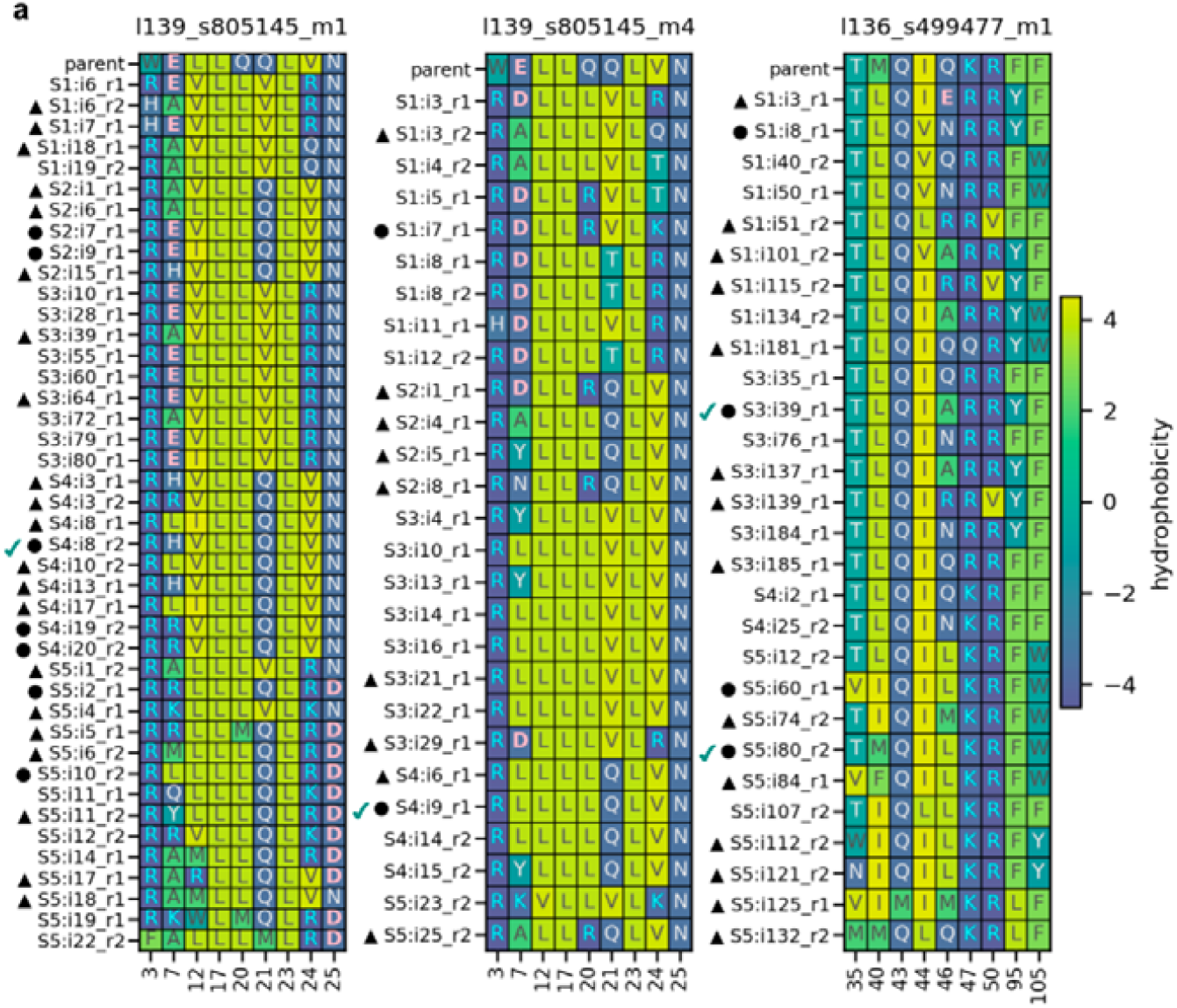
Interface sequence alignment of FEP-passing KRAS G12V–HLA-A*03:01 binders compared with their parent and sibling mutants. Residue boxes are colored by GRAVY hydrophobicity (see colorbar), with residue letters colored by type (negative: pink; positive: cyan; polar: light gray; hydrophobic: dark gray). Candidates passing MD but failing FEP are marked with triangles, while those passing both MD and FEP are marked with circles. SPR-validated binders are marked with ticks. Binder IDs use the following notation: l = length, s = seed, m = MPNN sequence, S = strategy, i = interface-optimized sequence, r = run of interface optimization.

**Supplementary Fig 2.**
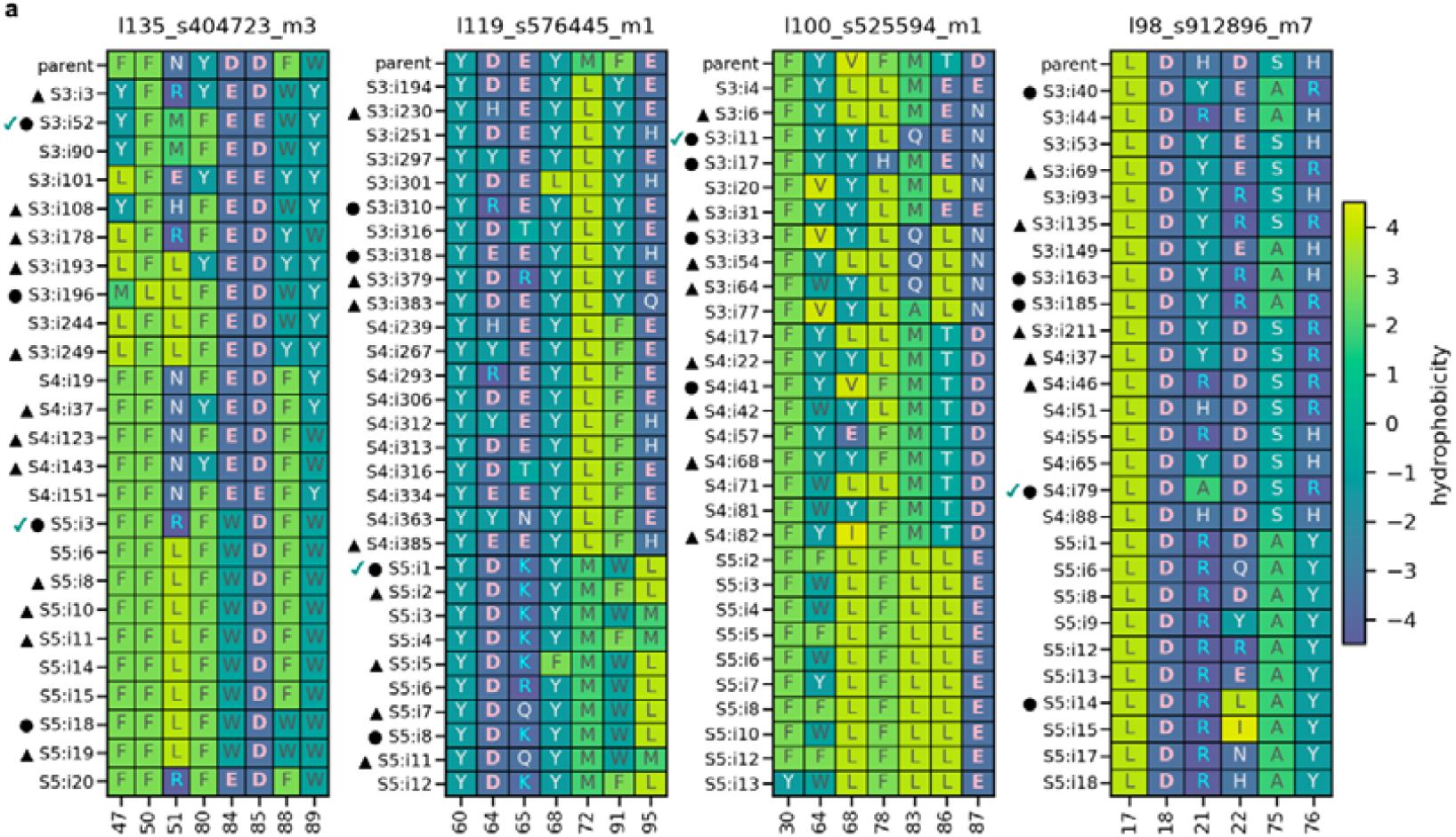
Interface sequence alignment of FEP-passing p53 R175H–HLA-A*02:01 binders compared with their parent and sibling mutants. Residue boxes are colored by GRAVY hydrophobicity (see colorbar), with residue letters colored by type (negative: pink; positive: cyan; polar: light gray; hydrophobic: dark gray). Candidates passing MD but failing FEP are marked with triangles, while those passing both MD and FEP are marked with circles. SPR-validated binders are marked with ticks. Binder IDs use the following notation: l = length, s = seed, m = MPNN sequence, S = strategy, i = interface-optimized sequence. The interface optimization run number was set to 1.

**Supplementary Fig 3.**
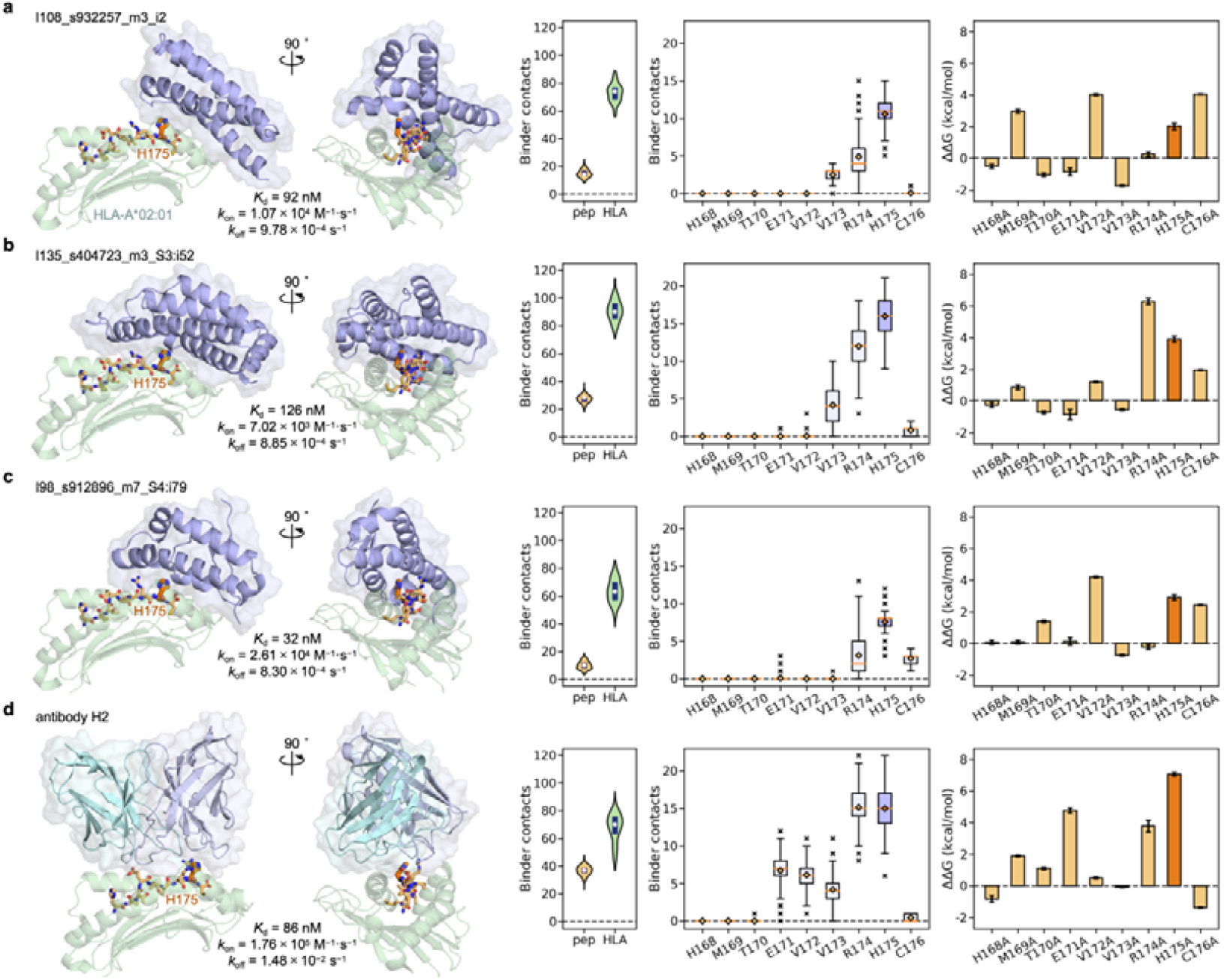
Comparison of designed binders with antibody H2 for p53 R175H–HLA-A*02:01. Structural views of p53 R175H–HLA-A*02:01 in complex with binders l108_s932257_m3_i2 **(a)**, l135_s404723_m3_S3:i52 **(b)**, l98_s912896_m7_S4:i79 **(c)**, or antibody H2 **(d)**. For each complex, the dissociation constant (*K*_d_), association rate (*k*_on_), and dissociation rate (*k*_off_) are reported alongside contact analysis and alanine-scanning FEP. Heavy atom contacts within 4 Å between binders and the neoantigen, HLA-A*02:01, and individual neoantigen residues were quantified during MD simulations. Antibody H2 affinities were measured using single-cycle kinetics, whereas binders were assessed using multi-cycle kinetics; because both employed the same ligand orientation (biotinylated pHLA on SA) and a standard 1:1 Langmuir model, the apparent *K*_d_ values are directly comparable, though minor assay-dependent differences may remain.

## Notes

### Competing Interest Statement

The authors have declared no competing interest.

